# Exploring Temporal and Sex-Linked Dysregulation in Alzheimer’s Disease Phospho-Proteome

**DOI:** 10.1101/2023.08.15.553056

**Authors:** Serhan Yılmaz, Filipa Blasco Tavares Pereira Lopes, Daniela Schlatzer, Rihua Wang, Xin Qi, Mehmet Koyutürk, Mark R. Chance

## Abstract

This study aims to characterize dysregulation of phosphorylation for the 5XFAD mouse model of Alzheimer’s disease (AD). Employing global phosphoproteome measurements, we analyze temporal (3, 6, 9 months) and sex-dependent effects on mouse hippocampus tissue to unveil molecular signatures associated with AD initiation and progression. Our results indicate 1.9 to 4.4 times higher phosphorylation prevalence compared to protein expression across all time points, with approximately 4.5 times greater prevalence in females compared to males at 3 and 9 months. Moreover, our findings reveal consistent phosphorylation of known AD biomarkers APOE and GFAP in 5XFAD mice, alongside novel candidates BIG3, CLCN6 and STX7, suggesting their potential as biomarkers for AD pathology. In addition, we identify PDK1 as a significantly dysregulated kinase at 9 months in females, and the regulation of gap junction activity as a key pathway associated with Alzheimer’s disease across all time points. AD-Xplorer, the interactive browser of our dataset, enables exploration of AD-related changes in phosphorylation, protein expression, kinase activities, and pathways. AD-Xplorer aids in biomarker discovery and therapeutic target identification, emphasizing temporal and sex-specific nature of significant phosphoproteomic signatures. Available at: https://yilmazs.shinyapps.io/ADXplorer

**Highlights:**

- Phosphorylation-level dysregulation surpasses protein expression
- Higher phospho-dysregulation in females, starting as early as 3-month time point
- Novel candidates BIG3, CLCN6, and STX7 exhibit consistent phospho-dysregulation
- Developed AD-Xplorer: Online tool to explore Alzheimer’s disease phospho-proteome

**In Brief:** This study investigates dysregulation of phospho-proteome in an Alzheimer’s disease (AD) mouse model, identifying consistent phosphorylation of established AD biomarkers APOE and GFAP, along with novel candidate biomarkers BIG3, CLCN6, and STX7. In addition, the study observes significant PDK1 dysregulation at 9 months, particularly in females. AD-Xplorer, our interactive tool for exploring temporal and sex-linked phosphorylation changes, protein expression, kinase activities, and pathway enrichment, empowers researchers to gain deeper insights into AD mechanisms and uncover novel biomarkers and therapeutic targets.

## 1. Introduction

Alzheimer’s disease (AD), a major cause of dementia, exhibits a high degree of heterogeneity and complexity in terms of risk factors, progression, and response to treatment [GBD, 2016; Ferreira, 2018; van der Flier, 2016; Devi, 2016]. However, the conventional postmortem diagnosis of AD based on histopathological examination of amyloid-beta (Aβ) has limitations, as abnormal Aβ deposition occurs before neurodegeneration and cognitive decline [Troncoso, 1996; Troncoso 1998]. Despite these limitations, most proteome-wide investigations in the literature have focused on postmortem brain samples, which cannot capture the dynamic molecular changes occurring during the progression of AD [Ferreira, 2018; van der Flier, 2016]. Elucidating the molecular mechanisms underlying abnormal Aβ deposition, as well as the timing and arrangement of the initiating and propagating pathways, could provide valuable insights into biomarkers and potential intervention strategies for predicting and managing AD progression in individual patients [Knopman, 2013].

The regulation of AD progression occurs at various levels, including DNA, RNA, and protein. The intricate nature of AD regulation is exemplified by transcriptomic and proteomic studies, which have shown a correlation of 0.45 between mRNA and protein levels [Sharma, 2015]. This suggests that these omics data are not individually sufficient to fully understand the important signaling events in Alzheimer’s Disease. Besides translational control, post-translational modifications (PTMs), such as phosphorylation, also play a crucial role in determining protein levels and variations [Needham 2019]. While abnormal protein phosphorylation in AD has been well documented [Kovacech, 2009] and Cdk5-tau hyperphosphorylation axis is a hallmark of AD [Noble, 2003], there is a lack of comprehensive studies investigating PTMs at a proteome-wide scale in brain tissue. Therefore, the exploration of phosphoproteomics holds significant potential in providing additional insights and identifying further pathways implicated in the progression of Alzheimer’s Disease [Marttinen, 2019].

Mouse models offer an excellent opportunity to study the temporal aspects of AD progression. For instance, a longitudinal study using a well-established amyloid-driven AD model (5XFAD mice) analyzed the perfusates from the hippocampus and revealed dysregulation in glucose and lipid metabolism prior to AD pathogenesis, demonstrating the feasibility of identifying early molecular events in disease progression [Gurel, 2019]. Our previous study [Lopes, 2022] utilized the same set of 5XFAD mice to better understand the temporal variation in protein expression in mouse brain for wild-type (WT) compared to the AD mouse model at various time points relevant to the key hallmarks of disease progression in humans.

In this study, we expand on our previous work [Lopes, 2022] and characterize the phospho-proteomic changes in 5XFAD mice at different time points relevant to the key hallmarks of disease progression in humans. The 5XFAD mice express five familial AD mutations (amyloid precursor protein and presenilin 1 genes), leading to the accumulation of amyloidogenic Aβ42 and a cascade of pathological changes that closely resemble human AD. Notably, these mice exhibit a significant increase in Aβ42 levels, resulting in the early onset of plaque deposition (around 2 months of age) that progressively spreads throughout the brain along with astrocytosis and microgliosis (around 4 months of age). Subsequently, neuronal loss occurs in the cortex and subiculum around 9 months of age. It is worth mentioning that, unlike the known human pathology, the Aβ plaques in the 5XFAD mouse model trigger the formation of neuritic tau plaque aggregates rather than neurofibrillary tangles, which are a crucial neuropathological hallmark of AD [Braunstein, 2016; Narasimhan, 2016]. Nevertheless, due to the overall similarities observed with human pathology, we designed a comprehensive study to explore the temporal and sex-related variations in protein phosphorylation, aiming to uncover new insights into known biomarkers and potentially identify novel targets for AD.

## 2. Experimental Procedures

### 2.1. Experimental Design

The study aimed to monitor the global phosphoproteome changes in the hippocampus of 5XFAD mice throughout different stages of Alzheimer’s Disease progression. To achieve this, label-free liquid chromatography-tandem mass spectrometry (LC-MS/MS) was employed. A total of 48 samples were analyzed, comprising sets of 16 male and female 5XFAD mice, as well as wild-type (WT) mice, which were run contemporaneously. The hippocampi were collected and processed at three time points: 3 months, 6 months, and 9 months. Each time point consisted of 8 samples for 5XFAD mice and 8 samples for WT mice. Furthermore, within each group, the samples were subdivided based on sex, resulting in 4 samples per subgroup. This comprehensive approach allowed for the investigation of the phosphoproteome changes in the hippocampus of male and female 5XFAD mice and male and female WT mice at each time point. All animal experiments were conducted in accordance with protocols approved by the Institutional Animal Care and Use Committee of Case Western Reserve University and performed according to the National Institutes of Health Guide for the Care and Use of Laboratory Animals.

In our study, we conducted several analyses to gain a comprehensive understanding of the phosphoproteome changes in the hippocampus of 5XFAD mice during the progression of Alzheimer’s Disease. First, we characterized phosphorylation patterns and compared them to protein expression levels to identify any correlations and to assess the complementarity between these two measures. Next, we investigated the temporal and sex-linked patterns in phosphorylation with a statistical analysis to identify specific dysregulated phosphopeptides between the WT and 5XFAD mice groups, revealing distinct molecular signatures associated with Alzheimer’s Disease progression. Moreover, we performed a biomarker analysis, identifying proteins that are consistently phosphorylated across different time points that could potentially serve as markers for Alzheimer’s Disease. For these identified proteins, we performed a computational prevalence analysis to quantify the stoichiometry of phosphorylation. In addition, we conducted kinase inference analysis to gain insights into the regulatory mechanisms involved in phosphorylation events. Finally, pathway analysis was conducted to identify the biological pathways and networks impacted by the observed phosphoproteome changes. Together, these analyses provided a multi-faceted approach to uncovering the complex dynamics of phosphorylation and its implications in Alzheimer’s Disease progression.

We validated our findings using independent validation data with a smaller number of samples, applying the same quantification method and statistical pipeline described.

### 2.2. Label-free quantitative proteomics and phosphoproteomics

Hippocampus tissue samples were collected from 5XFAD and wild-type mice at three different time points (three, six, and nine months). The samples were lysed using a 2% SDS solution supplemented with a protease inhibitor cocktail (Cat#P2714, Sigma) and PhosphoSTOP (Cat#04906837001, Roche). The protein concentration in the lysates was determined using the Bio-Rad Protein Assay Kit – BSA (Cat#5000002, Bio-Rad).

To process the lysates, we followed the FASP protocol (Wiśniewski 2009) and performed digestion using dual LysC (Cat#125-05061, Wako) and trypsin (Cat#90057, Thermo Fisher) endoproteinase. The digested samples were then desalted using C18 cartridges. For phosphopeptide enrichment, TiO2 enrichment spin tips were used. Samples were normalized to 300 ng of digest, and a blind and randomized approach was employed for LC-MS/MS acquisition. We used a Waters NanoAcquity UPLC chromatography system coupled to a Thermo Scientific Orbitrap Elite mass spectrometer (Thermo Fisher Scientific, CA).

The raw LC-MS/MS data was processed using Peaks v10.0 Software (Bioinformatics Solutions, ON, CA). Peptide identification was performed within Peaks using the UNIPROT database (UNIPROT_MOUSE_091219, # of entries=17026). The PEAKS search parameters included a mass error tolerance of 10 ppm for precursor ions, a mass tolerance of 0.6Da for fragment ions, trypsin enzyme specificity, and fixed variations of carbamidomethylation, as well as variable modifications of methionine oxidation and phosphorylation at serine, threonine, and tyrosine residues. The search allowed for one missed cleavage. Label-free phosphopeptide identification followed the default target decoy approach, with a PEAKS peptide score (−10logP) threshold of ≥ 15 and an FDR threshold of 1%. The abundance of individual peptides was determined by calculating the area under the curve (AUC). Supplementary Data 1 contains the output excel files for all three time points, along with the corresponding sample information.

### 2.3. Statistical Rationale

To ensure data quality, we performed a number of additional preprocessing steps. First, we filtered the peptides for monophosphorylated peptides. Next, we excluded quantifications with small intensities (<10^4) from the analysis by treating them as missing values. Additionally, we filtered out peptides that have numerous missing values. Specifically, a peptide was included in the analysis only if it is identified in a minimum of 3 samples for both the WT and 5XFAD groups. We also filtered out 2 samples in the 9 month time point (one female and one male sample in the WT group) that exhibited substantially low mean intensities across all peptides (Supp Fig 1). In addition, to alleviate potential imbalances between samples, we applied a preprocessing step that centers the log2 transformed intensities of all samples. This process ensures that the mean value across all peptides is the same for every sample, irrespective of their WT/5XFAD group. We investigated the differences between 5XFAD and WT samples by performing a moderated t-test, which utilizes an empirical Bayes method to shrink the sample variances towards a common value and to augment the degrees of freedom for the individual variances [Smyth, 2004].

To identify potential phosphorylation-based biomarkers that are consistent across time points, we conducted an analysis at the protein level. This approach was chosen due to the high amount of missingness at the phosphopeptide level, particularly when comparing different time point experiments (e.g., 3 months, 6 months, 9 months). The rate of missingness at the protein level was considerably less compared to the number of phosphopeptides identified across all three time points. By focusing the analysis at the protein level, we aimed to address this challenge and facilitate the assessment of consistency across various time points or groups. For this purpose, we computed the mean log-fold changes at the protein level and used a moderated t-test with the Satterthwaite approximation [Satterthwaite, 1946] to compare 5XFAD and WT groups (see supplement for details). To assess consistency across time points, we introduced a simple statistic called the consistency score, which combined the log2-fold change results of individual time points. This score represents the total log2-fold change across time points, considering any missing values as log2-fold change of 0. This approach allowed us to account for changes in the same direction while canceling out those with opposite directions, facilitating the identification of proteins with consistent phosphorylation patterns.

To quantify the stoichiometry of phosphorylation, we conducted a computational analysis, calculating the ratio of the top phospho-peptide to the top unmodified peptide with highest intensities for each protein, separately for each sample in both 5XFAD and control groups. For this purpose, given the technological constraints in directly measuring small phosphorylation intensities in unenriched data [Lopes, 2022], we used the phospho-enriched data to derive a scaling factor for each sample. This scaling factor enabled us to estimate the phosphorylation intensities in the unenriched data using the phospho-enriched data. To achieve this, we first identified common proteins between the two datasets and determined the highest phospho-peptide intensity for each protein. Then, we assessed how many times, on average, the intensities in the phospho-enriched data was higher compared to the unenriched data for each sample using the geometric mean (Supp. Fig 2). By dividing the intensities in the phospho-enriched data by this scaling factor, we obtained the estimated phosphorylation intensities for each protein in the unenriched samples. Finally, we calculated the prevalence of phosphorylation for each protein and sample, reporting mean, minimum and maximum values across samples in both AD and control groups.

For the kinase analysis, we considered kinases with at least two known targets identified in our dataset and performed the kinase activity inference using the RoKAI algorithm, which makes use of a functional network to improve the accuracy and robustness of the inference [Yılmaz, 2021]. To identify biological pathways and networks impacted by the observed phosphoproteome changes, we performed a quantitative pathway enrichment analysis based on the mean phosphorylation (log2-FC) of proteins. This approach helped us overcome the limitations of relying on arbitrary significance thresholds and provided a more interpretable measure of pathway enrichment, enabling us to quantify the magnitude of enrichment in addition to identifying the significant pathways affected by the observed changes in the phosphoproteome. We conducted the pathway analysis using Reactome pathways [Jassal, 2020] that met the following inclusion criteria. We excluded pathways with a large portion of missing data, considering only those with at least two proteins identified in our dataset. Additionally, we required that these proteins accounted for at least 10% of all proteins in the pathway. To avoid redundancy, we filtered out highly similar pathways sharing the same protein sets, retaining the smaller pathway in cases of duplication.

In our phosphopeptide analysis, to determine phosphopeptides that are of significance, we applied two sets of thresholds. The first set, which includes a p-value cutoff of p ≤ 0.1 followed by a fold change cutoff of 0.5 ≥ FC ≥ 2, enables us to compare results across different time points and/or different sexes. This simple criteria provides a consistent framework for making comparisons between different groups. However, to address the multiple comparisons issue and to enable individual discoveries, we also employed a second set of thresholds. This involves utilizing the Benjamini-Hochberg procedure to apply a false discovery rate (FDR) cutoff of less than 0.1, ensuring statistically significant findings that are unlikely to be due to random noise or false positives.

Lastly, to aid in the interpretation of our findings, we used the interactive data browser feature of RokaiXplorer application^1^ to develop the online tool AD-Xplorer^2^. In this tool, the users can interactively explore the AD phospho-proteomic dataset that we present in this paper, as well as the proteomic dataset that is a result of our previous study on the same set of mice [Lopes, 2022]. In addition to the previous analyses, AD-Xplorer includes a gene ontology (GO) term enrichment performed with an over-representation analysis (chi-squared test with Yates’ correction) based on a screened set of peptides or proteins (default cutoffs: p ≤ 0.1 and 0.5 ≥ FC ≥ 2).

## 3. Results

### 3.1. Exploring Complementarity: Dysregulation in Phosphorylation vs. Protein Expression

We started our investigation by analyzing the trends in peptides identified through phosphorylation, and comparing them with peptides identified from protein expression data of a previous study on the same set of mice [Lopes, 2022]. To ensure consistency, we applied the same thresholds that were used in the prior protein expression analysis. Specifically, we used 0.5 ≥ FC ≥ 2 and p ≤ 0.1 thresholds to screen phospho-peptides for differential phosphorylation, comparing 5XFAD mice with WT mice. Our initial observations revealed a gradual increase in peptides identified based on protein expression as time progressed (Supp Fig. 3). Similar trends were observed in the number of peptides identified through phosphorylation, with a slight decrease at the 6-month time point. Notably, although protein expression yielded a higher number of peptides that pass the cutoffs (around 2-5 times more), this is mainly due to higher numbers of peptides in the protein expression data (around 4-8 times more). However, when considering the percentage of peptides that satisfy the screening criteria relative to all peptides, phosphorylation exhibited higher prevalence compared to protein expression. In mixed-sex analysis, phosphorylation resulted in 2.26%, 1.96%, and 2.56% of peptides identified for 3, 6, and 9-month time points, while protein expression resulted in 0.51%, 0.73%, and 1.36%. This corresponds to 4.4x, 2.7x, and 1.9x higher phosphorylation prevalence compared to protein expression.

Additionally, our sex-specific analysis showed a notably higher percentage of differentially phosphorylated peptides that pass the cutoffs in females for 3 and 9 months (6.67% and 6.56% respectively), and 1.85% for 6 months. These percentages correspond to 4.4x, 1.7x, 5.2x higher prevalence of phosphorylation in females compared to protein expression, and 4.7x, 1.0x, 4.4x higher prevalence compared to males, respectively for 3, 6, and 9 month time points. These findings suggest that phosphorylation undergoes significant changes in the 5XFAD hippocampus at an early stage, potentially surpassing protein expression changes in magnitude.

Next, to assess the complementarity of phosphorylation information to protein expression, we examined their correlation at each time point. Comparing phosphorylation at the peptide-level and protein expression, we observed only a minor correlation (R^2=0.09, 0.02, and 0.12 for 3, 6, and 9 months; Supp Fig. 4). However, when comparing protein expression with phosphorylation at the protein-level (mean phosphorylation), we found a higher, yet still modest, correlation for all time points (R^2=0.20, 0.07, and 0.27 for 3, 6, and 9 months). On the other hand, as a reference point, investigating the correlation between phosphorylation at the phosphosite level and phosphorylation at the protein level (mean phosphorylation), we observed a much stronger correlation (R^2=0.56, 0.44, and 0.56 for 3, 6, and 9 months). These findings highlight that changes in phosphorylation and protein expression are not redundant but rather complementary to each other, as the lack of a strong correlation indicates they both provide distinct and valuable information.

### 3.2. Temporal Dynamics: Phosphorylation Profile of 5XFAD Mouse Model Over Time

After the filtering and quality control steps, we identify 3287, 3198, and 2748 phosphopeptides in the 3-month, 6-month, and 9-month datasets respectively, all contributing to a total of hippocampal coverage of 1782 phosphorylated proteins. The volcano plots for each dataset highlight a dramatic accumulation of differentially phosphorylated peptides at the 6-Month time (screening threshold: 5XFAD/WT 0.5 ≥ FC ≥ 2 and p ≤ 0.1; Fig. 1, A–C). Respectively in the 3, 6, and 9 month time points, 48, 28, and 30 phosphopeptides were upregulated while 25, 34, and 40 phosphopeptides were downregulated. Thus, half of these differentially phosphorylated peptides are upregulated in the 5XFAD mice.

**Figure 1.**
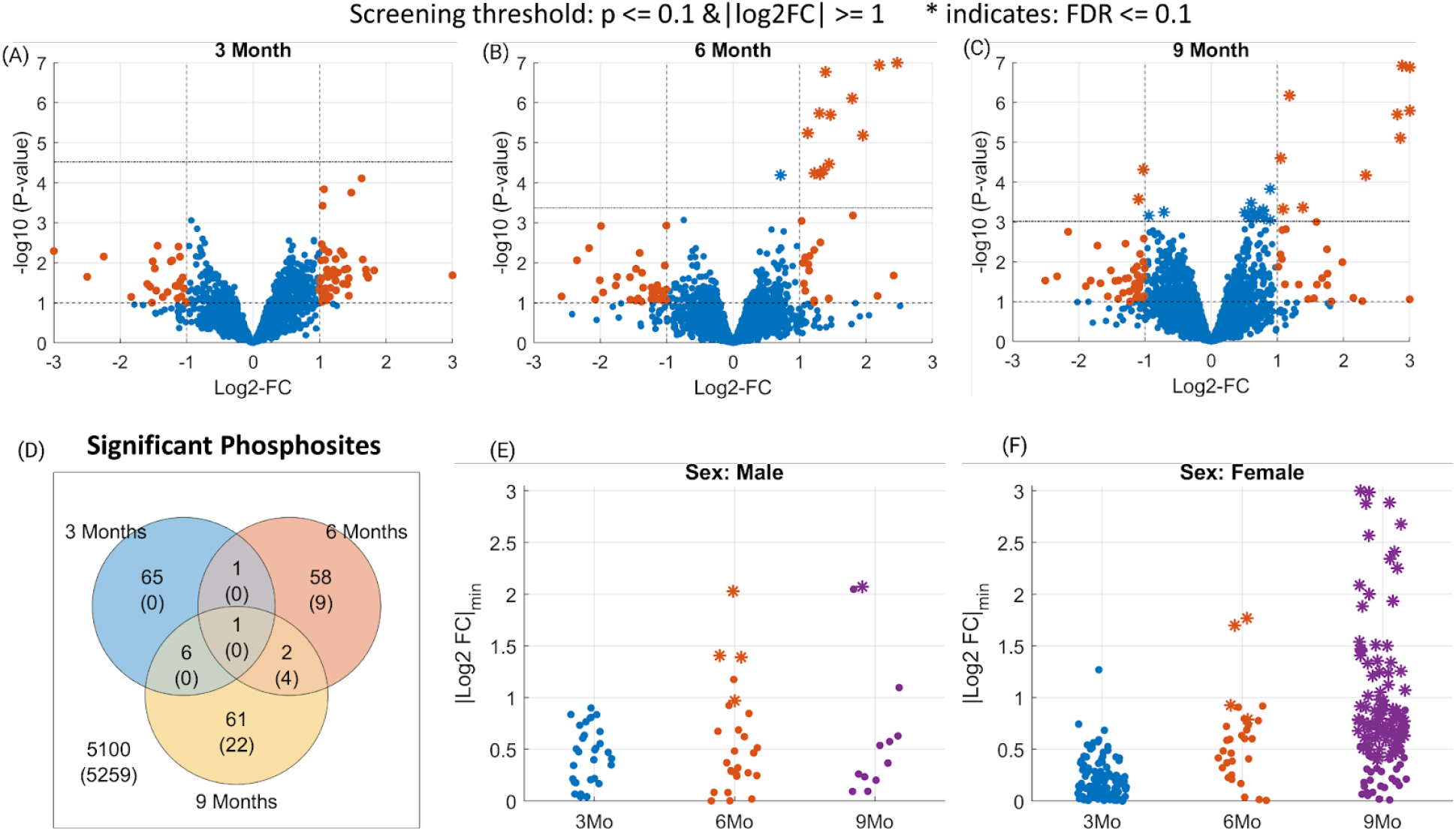
Phosphosites identified as differentially phosphorylated for Alzheimer’s disease at different time points. (Panels A-C) Volcano plots indicating the screened phosphosites with high or low phosphorylation at 3/6/9 month time points. The dashed horizontal lines in each plot indicate the statistical significance threshold (p<0.1 and FDR ≤ 0.1) and the vertical line the threshold on fold changes (>2). The statistically significant phosphosites at FDR ≤ 0.1 level are marked as stars (*). (Panel D) Venn diagram showing the number of phosphosites that pass the screening (p ≤ 0.1 and 0.5 ≥ FC ≥ 2) for 3/6/9 months data. Whereas, the numbers in parenthesis indicate the significant ones at FDR ≤ 0.1 level. (Panels E-F) Identified phosphosites that pass the screening in sex-specific analysis. The y-axis indicates a lower-bound of the 95% confidence intervals for each phosphosite.

To characterize the temporal persistence of these phosphorylation signatures, we analyzed the overlap among the three time points. The Venn diagram highlights the transitory nature of phosphorylation signaling in Alzheimer’s disease, with less than 5% of all phosphopeptides being dysregulated in two or more timepoints (Fig. 1, D). Indeed, only glial fibrillary acidic protein (GFAP) passed the screening thresholds across all three time points, exhibiting consistent hyperphosphorylation across time. This complements the proteome changes of the same set of mice that found GFAP protein expression to be significantly upregulated at all time-points [Lopes, 2022]. This is expected as GFAP is a glial and endothelial cell phenotypic marker and doubles as a classical astrogliosis marker [Pixley, 1984; Levin, 2009], which in turn plays a central role in AD neuroinflammation [Bilbao, 2014; McColl, 2018].

Sex differences in frequency, disease burden, and risk factors for AD are major long-standing questions in the field. Here, we exploited our sex-balanced experimental design to characterize the sex-dependent variations in phosphorylation across Alzheimer’s disease progression. We identified the differentially phosphorylated peptides separately for female and male groups at each time point (5XFAD male vs. WT male, and 5XFAD female vs. WT female, p≤0.1 and 0.5≥FC≥2). Male mice exhibit a consistent phosphopeptide signature throughout all timepoints, respectively 33, 38, and 23 differentially phosphorylated peptides at 3, 6, and 9 months (Fig. 1, E). In contrast, female 5XFAD mice display a distinct phosphopeptide profile with 169, 36, and 126 phosphopeptides at 3, 6, and 9 months respectively (Fig. 1, F). In total, the number of differentially phosphorylated peptides identified for female mice is more than three times greater than that for male mice (Supp. Fig. 5).

### 3.3 Temporal and Sex-Specific Analysis: Phosphopeptide Discoveries in 5XFAD Mouse Model

To control for false discoveries, we employed a moderated t-test and applied the Benjamini-Hochberg procedure with a false discovery rate (FDR) threshold of 0.1. We imposed this significance threshold to identify individual phosphopeptides that showed significant dysregulation in each of the 3/6/9 month time points separately. In Figure 1, the phosphopeptides that met this significance threshold (FDR ≤ 0.1) are denoted by asterisks in panels A to C. Our analysis revealed an increasing number of dysregulated phosphopeptides identified at 3, 6, and 9 months, specifically 0, 13, and 26, respectively (Figure 1, D). Among these dysregulated phosphopeptides, four were found to be common between 6 and 9 month time points: Apolipoprotein E (APOE)-T140, Brefeldin A-inhibited guanine nucleotide-exchange protein 3 (BIG3)-S1646, Syntaxin 7 (STX7)-T79, and GFAP-S12.

Next, among the phosphopeptides that we identify (FDR ≤ 0.1), we investigated which of these also met the fold change threshold (5XFAD/WT 0.5 ≥ FC ≥ 2). We observed a total of 12 phosphopeptides for the 6-month time point and 12 phosphopeptides for the 9-month time point. Notable ones among these include APOE-S139, Chloride voltage-gated channel 6 (CLCN6)-S726 and S685, and GFAP-T40, which exhibit significant upregulation exclusively in the 6 month time point. Whereas, CLCN6-S774, STX7-S45, GFAP-Y13, and Amyloid-beta precursor protein (A4)-S441 are among phosphopeptides exhibiting significant upregulation only in the 9 month time point. Overall, we observe 2 phosphopeptides that are identified as significant in both 6 and 9 month time points, which are BIG3-S1646 and GFAP-S12. Apart from the known AD hallmarks, we detect the upregulation of STX7 and CLCN6, which implies the involvement of AD in the dysregulation of the late endocytic pathway [Jentsch, 2018]. For a comprehensive list of the phosphopeptides identified as significant in this analysis, please refer to Supplementary Data 2.

When we explore the sex-specific differences, we observe respectively 0, 4, 142 phosphopeptides for females and 0, 4, 1 phosphopeptides for males that pass the significance threshold (FDR ≤ 0.1) for 3, 6, and 9-month data. At 6 months, APOE-T140, APOE-S139, CLCN6-S685, BIG3-S1646 are the discoveries for the female group, whereas APOE-T140, Presenilin-1 (PSN1)-S353, Brevican core protein (PGCB)-T544, GFAP-S12 are for the male group. In females, there are 2 phosphopeptides that are identified as significant in both 6 and 9 month time points, which are BIG3-S1646 and APOE-T140. Whereas, in males, only APOE-T140 overlaps between 6 and 9 month time points.

Next, to further filter the 142 phosphopeptides we identified in the 9 month female group (FDR≤0.1) and uncover sites of biological interest, we investigated those that exhibit a large difference in fold changes (5XFAD/WT 0.5 ≥ FC ≥ 2). We observed 88 out of these 142 findings also passed this fold change threshold. Among these, the peptides that exhibit larger differences in fold changes (0.25 ≥ FC ≥ 4) are: BIG3-S1646, GFAP-Y13, A4-S441, Myelin-associated glycoprotein (MAG)-S547, Microtubule-associated protein 2 (MTAP2)-S739, Vacuolar protein sorting-associated protein 53 homolog (VPS53)-S377, Oxysterol-binding protein 1 (OSBP1)-S349, cGMP-dependent 3’,5’-cyclic phosphodiesterase (PDE2A)-S911, and Microtubule-associated protein 6 (MAP6)-S905. A STRING network analysis of these top phosphorylated proteins reveals an enrichment for Golgi apparatus proteins (data not shown). This suggests that a dysregulation of the Golgi apparatus in female mice might contribute to the higher disease burden in female mice especially in late AD stages.

To identify phosphopeptides that are differentially regulated between males and females, we compare 5XFAD/WT fold changes in females vs. the fold changes in males. We identify 4 phosphopeptides that are significantly different in the 9 month time point (FDR≤0.1), these are MTAP2-S739, Adhesion G protein-coupled receptor B2 (AGRB2)-S1277, 14-3-3 protein epsilon (1433E)-S210, and BTB/POZ domain-containing protein KCTD12 (KCD12)-S178. Next, we investigate phosphopeptides that were significant in females, but not in males (FDR≤0.1), that also exhibit a large fold change difference between females and males (0.25≥FC≥4). Based on this analysis, we identify 23 peptides. Notable ones among these include: BIG3-S1646, OSBP1-S349, VPS53-S377, vacuolar protein sorting-associated protein 35 (VPS35)-S786, MAG-S547, EH domain-containing protein 3 (EHD3)-S456, receptor-type tyrosine-protein phosphatase zeta (PRPTZ)-S576, synapsin-1 (SYN1)-S67, NMDA 2B glutamate receptor ionotropic (NMDE2)-S948, and 14-3-3 protein zeta/delta (1433Z)-S207. We provide the rest of the significant findings in Supplementary Data 2 and they can also be browsed from the AD-Xplorer application.

### 3.4. Potential Alzheimer’s Disease Markers: Consistent Phospho-Dysregulation Across Time

To identify potential biomarkers of disease progression, we conducted a longitudinal analysis of phosphoproteome profiling. We examined the dysregulation of proteins across all time points and calculated a consistency score based on their mean log2 fold change at the protein level and the total fold change across the three time points. Based on this analysis, we identified 31 proteins that were consistently upregulated and 18 proteins that were consistently downregulated, all at a false discovery rate (FDR) of 0.1 or lower. Among these proteins, several proteins emerged as candidate biomarkers. The nine proteins exhibiting consistently increased phosphorylation levels are: APOE, GFAP, BIG3, CLCN6, Protein tyrosine phosphatase receptor type C (PTPRC), A4, STX7, Gap junction alpha-1 protein (GJA1), Plexin domain containing protein 2 (PXDC2). On the other hand, three proteins exhibiting consistently decreased phosphorylation are: Disks large homolog 4 (DLG4), Rad and gem-like GTP-Binding protein 2 (REM2), and general receptor for phosphoinositides 1-associated scaffold protein (GRASP) - see Fig. 2.

**Figure 2.**
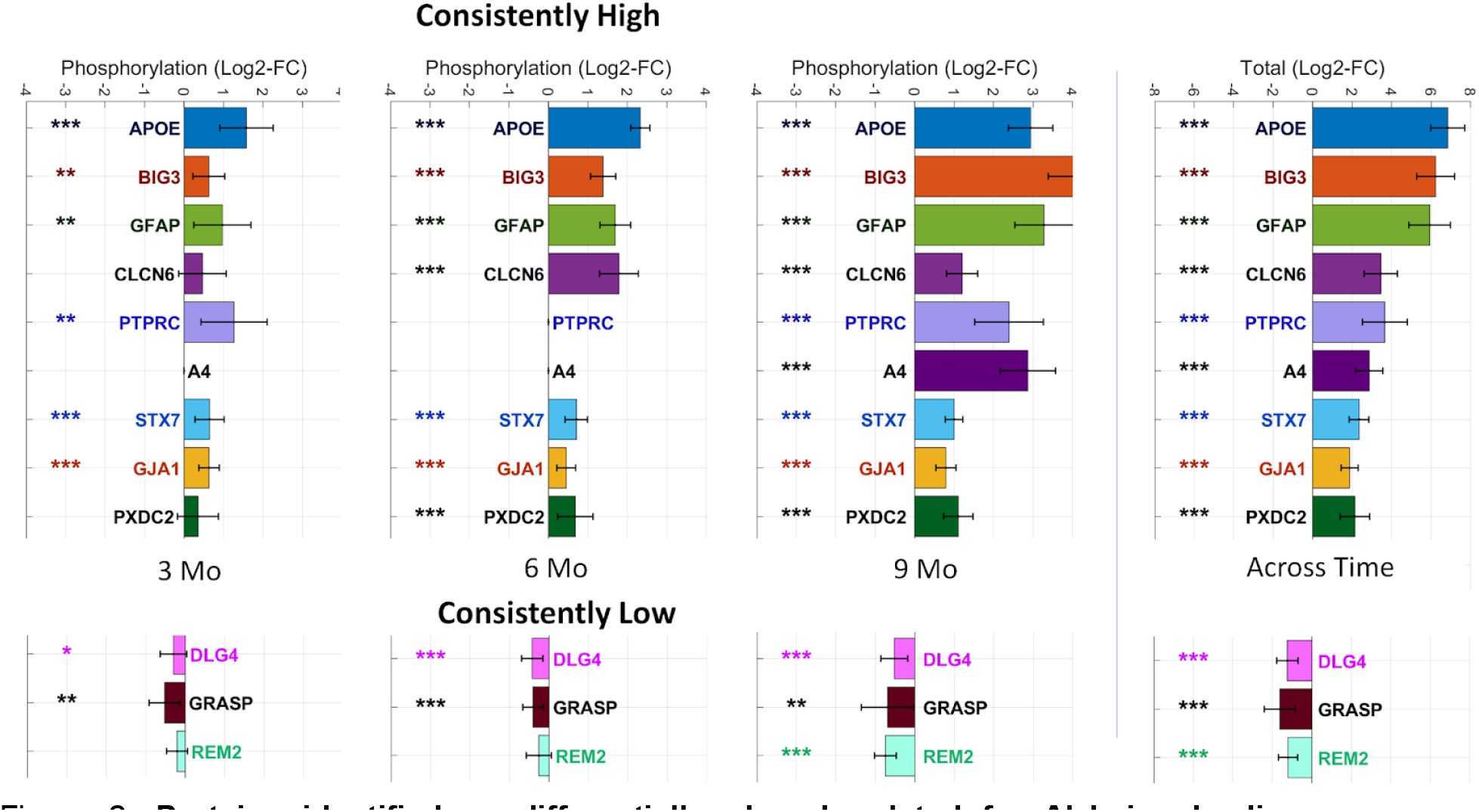
Proteins identified as differentially phosphorylated for Alzheimer’s disease across different time-points. (Left panel) The bars indicate the mean phosphorylation for each protein identified as consistently phosphorylated and the error bars indicate 95% confidence intervals. Proteins are marked according to their statistical significance levels: * for p ≤ 0.1, ** for FDR ≤ 0.1 and *** for FDR ≤ 0.01.

In our sex-specific analysis, we observe distinct patterns between females and males. Specifically, we found 41 consistently high and 42 consistently low proteins in females, compared to 11 consistently high and 1 consistently low proteins in males. Interestingly, we observe 10 proteins that are identified as consistent in both females and male specific analysis, including APOE, BIG3, CLCN6, GFAP, STX7, PSN1, GJA1, Serine incorporator 1 (SERC1), Actin-binding LIM protein 1 (ABLM1), and Brain acid soluble protein 1 (BASP1). Among these, only BASP1 exhibits decreased phosphorylation.

Moreover, we observe two proteins, Neural proliferation differentiation and control protein 1 (NPDC1) and Ubiquitin-conjugating enzyme E2 variant 3 (UEVLD), that are identified as consistently hyper-phosphorylated in males but not in females. On the other hand, when we examine the proteins identified exclusively in female-specific analysis, we observe a large number of proteins that exhibit a considerable total fold change across the three time points (0.25 ≥ Total FC ≥ 4). These proteins include PXDC2, GRASP, MAG, Electrogenic sodium bicarbonate cotransporter (S4A4), DnaJ homolog subfamily C member 5 (DNJC5), Src substrate cortactin (SRC8), Synaptotagmin-2 (SYT2), GRIP1-associated protein 1 (GRAP1), SRSF protein kinase 2 (SRPK2), Syntaxin-binding protein 5 (STXB5), Galectin-1 (LEGL), Synaptophysin (SYPH), Basigin (BASI), and Peroxisomal biogenesis factor 19 (PEX19). Among these, only PXDC2, S4A4, SRC8, DNJC5, and GRASP are also identified as consistently phosphorylated in the mixed sex analysis. Overall, our analysis reveal 19 consistently high and 27 consistently low proteins that are specific to the female-specific analysis and not in the mixed sex or male-specific analyses. For more detailed findings, please see Supplementary Data 3.

### 3.5. Assessing Functional Significance: Stoichiometry of Candidate Phosphorylation Markers

To assess phosphorylation stoichiometry, we conducted a computational analysis, computing the ratio of the top phosphopeptide to the top unmodified peptide for each protein, separately for each sample in both AD and control groups. Due to the limited ability of available technology to directly measure small intensities in unenriched samples, we utilized the phospho-enriched data to establish a scaling factor for each sample, estimating the intensities in unenriched data. The average phospho-enrichment factor was approximately 500 times (range: 244-912; Supp. Figure 2). As a result of this analysis, we report the mean phosphorylation rate across AD samples, along with the corresponding minimum and maximum values for each protein (shown in brackets). Notably, APOE and GFAP showed minimal phosphorylation rate (<1%) across all samples and time points, calling into question the biological significance of these modifications. In contrast, STX7 and GJA1 exhibited considerable phosphorylation rate, particularly at 9 months (STX7: 26.2% [18.0%-43.5%]; GJA1: 22.8% [14.5%-38.4%] at 9 months). We also observed a high phosphorylation rate of BIG3 at 6 months (33.5% [26.1%-43.1%]), while it was not identified at 3 months or 9 months. Detailed results are provided in Supplementary Data 4.

### 3.6. Inferred Kinase Activities: Temporal and Sex-Specific Analysis

Phosphorylation, regulated by kinases, plays a crucial role in various cellular mechanisms and is of significant interest in Alzheimer’s Disease (AD) research. To understand the impact of phosphorylation on AD, we focused on characterizing kinase activity. Using RoKAI, a tool for inferring kinase activity, we identified the top kinases at each timepoint. We performed separate analyses for each sex (male and female) as well as a combined, mixed-sex analysis (Fig. 3).

**Figure 3.**
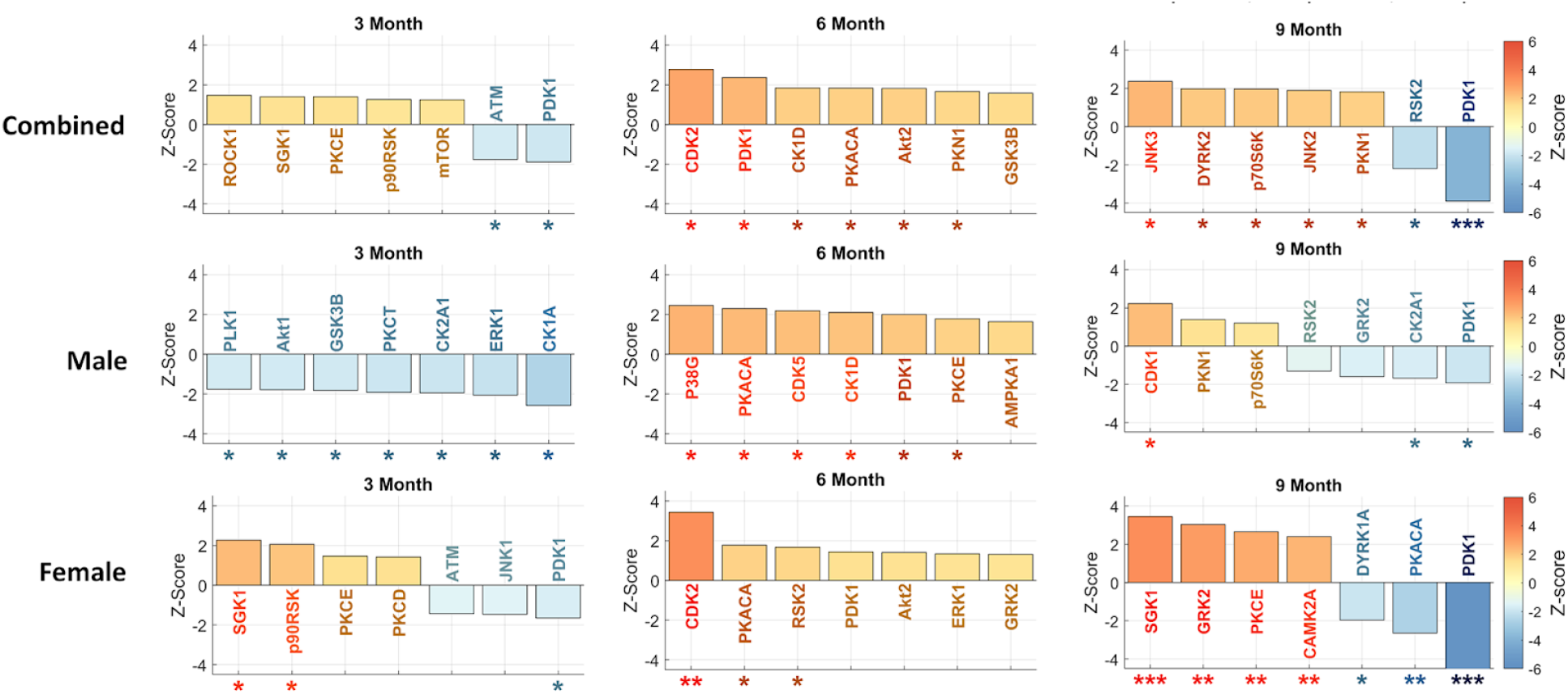
Inferred kinase activities across 3/6/9 months time-points and different sexes. For each subplot, the top seven most significant kinases are shown. The error bars indicate 95% confidence intervals. Kinases are marked according to their statistical significance levels: * for p ≤ 0.1, ** for FDR ≤ 0.1 and *** for FDR ≤ 0.01.

Investigating significant kinase dysregulation at FDR≤0.1 level in our mixed sex analysis, we observe that phosphoinositide-dependent kinase-1 (PDK1) exhibits downregulation at 9 months. In the sex-specific analysis, we find a higher level of dysregulation in females compared to males, identifying several dysregulated kinases in females but none in males at the FDR≤0.1 level. Specifically, we identify upregulation of cyclin-dependent kinase 2 (CDK2) in the 6-month female group, as well as upregulation of serum/glucocorticoid-regulated kinase 1 (SGK1), G protein-coupled receptor kinase 2 (GRK2), protein kinase C epsilon (PKCE), and calcium/calmodulin-dependent protein kinase 2 alpha (CAMK2A) in 9-month females. Additionally, we identify downregulation of PDK1 and protein kinase CAMP-Activated catalytic subunit alpha (PKACA) in the 9-month female group. These results underscore the distinctive roles of kinases in AD progression between female and male mice. For detailed results, please refer to Supplementary Data 5.

### 3.7. Dysregulation of PDK1: Phosphorylation, Protein Expression, and mRNA Analysis

In the 5XFAD mouse model, we observed downregulation of PDK1 activity at three and nine months, but a short-lived upregulation at the six-month time point. Specifically, we observe divergent patterns of PDK1 regulation between female and male 5XFAD mice. The male dataset demonstrated significant upregulation of PDK1 activity at six months (p-value ≤ 0.1), while the female dataset shows significant downregulation as early as the three-month time point and identifies PDK1 upregulation (p-value ≤ 0.1) as a top kinase (Fig. 3).

To further investigate the regulation of PDK1 at the protein expression level, we examined a previous study that analyzed global proteomic changes in the same set of 5XFAD mice (Fig. 4A) [Lopes, 2022]. The data revealed distinct expression patterns of PDK1 between male and female 5XFAD mice at six and nine months. Male mice showed downregulation of PDK1 at both timepoints, while female mice exhibited little change at six months followed by significant upregulation at nine months (Fig. 4A). These findings indicate that changes in protein expression and phosphorylation of PDK1 are not entirely concordant. Specifically, at six months, male mice showed a trend of PDK1 downregulation, but inferred PDK1 activity was significantly upregulated (p-value ≤ 0.1) (Fig. 3). At nine months, the upregulation of PDK1 expression in female mice was counteracted by a significant downregulation of inferred PDK1 activity (FDR≤0.01). Conversely, at nine months, male mice exhibited a downregulation trend in both PDK1 protein expression and inferred activity. These findings suggest that PDK1 is under complex regulation that changes throughout the progression of AD in both male and female mice.

**Figure 4.**
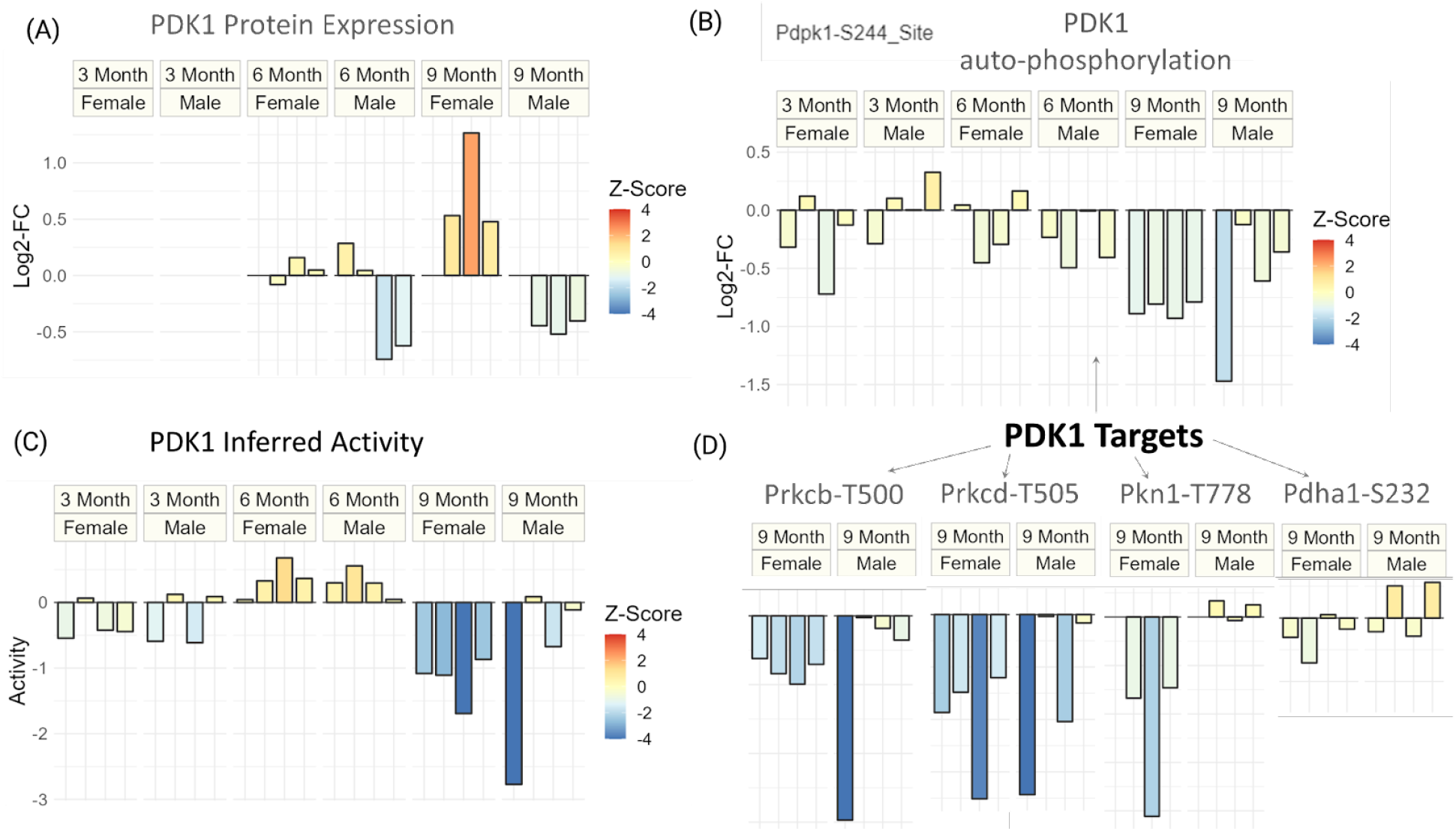
Dysregulation of PDK1 at protein expression and phosphorylation level for different sexes and timepoints of Alzheimer’s disease. (Panel A) Protein expression (log2-FC) of PDK1 for all timepoints and sexes. (Panel B) The auto-phosphorylation (log2-FC) of PDK1 for all timepoints and sexes. (Panel C) Inferred activity score of PDK1 for all timepoints and sexes. (Panel D) Phosphorylation (log2-FC) of known targets of PDK1 for 9 month data. For all plots, the coloring is done to reflect the degree of statistical significance based on z-scores.

To gain insights into the regulation of PDK1 at the transcriptional level, we examined a longitudinal 5XFAD transcriptomic study [Forner, 2021]. We found that PDK1 mRNA levels remained relatively stable with a fold-change (5XFAD/WT) from four to eighteen months (data not shown). These results indicate that the changes in PDK1 detected in 5XFAD mice are primarily regulated at the protein level and involve complex mechanisms.

To further investigate the regulation of PDK1 in AD mice, we analyzed the phosphorylation changes at PDK1’s constitutively activated site, S244 (Fig. 4, B). At three and nine months, we observed that inferred activity and autophosphorylation levels for PDK1 are concordant (Fig. 4, B-C). However, at six months, while PDK1 inferred activity was upregulated, phosphorylation at the autophosphorylation site was not, suggesting that constitutive activation of PDK1 alone does not explain its regulation at six months. We observe a significant downregulation of the PDK1-S244 site only at 9 months (p-value ≤ 0.1).

To explore the downstream signaling consequences of PDK1 at advanced stages of Alzheimer’s Disease, we examine the phosphorylation levels of PDK1’s known targets that are quantified in our dataset (Fig. 4D). We observe that the phosphorylation levels of protein kinase C beta (PRKCB) and protein kinase C delta (PRKCD) are particularly concordant with the inferred downregulation of PDK1 at nine months.

### 3.8. Interpreting Biological Significance: Pathway Analysis of Phosphoproteome Changes

To understand the relation between Alzheimer’s Disease (AD) and various pathways, we conducted an analysis of phosphoproteomic datasets at different time points and performed a quantitative enrichment analysis using Reactome pathways.

Based on this analysis, we identify 5, 16, and 67 pathways that were significantly enriched at 3, 6, and 9 months respectively (FDR≤0.1). While most of these pathways are specific to particular timepoints, the regulation of gap junction activity was consistently enriched across all timepoints. Additionally, the scavenging by Class A Receptors pathway showed significant enrichment at both 6 and 9 months (Table 1). Notable pathways unique to specific time points include: Cation-coupled Chloride cotransporters, RHO GTPases activate PAKs, DARPP-32 events, and RAF/MAP kinase cascade at 3 months; EPH-ephrin mediated repulsion of cells, noncanonical activation of NOTCH3, Nuclear signaling by ERBB4, NRIF signals cell death from the nucleus, and CS/DS degradation at 6 months; and GABA synthesis, release, reuptake and degradation, insertion of tail-anchored proteins into the endoplasmic reticulum membrane, MTOR signaling, norepinephrine neurotransmitter release cycle, and dopamine neurotransmitter release cycle at 9 months (Table 1).

**Table 1.** Reactome pathways identified as significant in AD phospho-proteome for 3/6/9 month time points. Each row corresponds to a different pathway. Up to seven significant pathways are shown for each time point. “# of proteins” column indicates the number of proteins in the pathway, the first number is for proteins identified in our data and the second one (after /) is for total number of proteins.

|  | Pathway | # of Proteins | Z-Score | FDR |
| --- | --- | --- | --- | --- |
| 3 Month | Regulation of gap junction activity | 2 / 3 | 4.60 | 0.0019 |
|  | Cation-coupled Chloride cotransporters | 2 / 7 | 4.13 | 0.0053 |
|  | RHO GTPases activate PAKs | 10 / 20 | 3.34 | 0.08 |
|  | DARPP-32 events | 7 / 16 | 3.31 | 0.08 |
|  | RAF/MAP kinase cascade | 26 / 91 | -4.28 | 0.004 |
| 6 Month | Regulation of gap junction activity | 2 / 3 | 3.59 | 0.021 |
|  | Scavenging by Class A Receptors | 2 / 7 | 8.64 | 0 |
|  | EPH-ephrin mediated repulsion of cells | 5 / 30 | 4.90 | 0.00021 |
|  | Noncanonical activation of NOTCH3 | 2 / 5 | 4.62 | 0.00054 |
|  | Nuclear signaling by ERBB4 | 2 / 9 | 4.21 | 0.0028 |
|  | NRIF signals cell death from the nucleus | 2 / 10 | 4.10 | 0.029 |
|  | CS/DS degradation | 2 / 7 | -3.37 | 0.0033 |
| 9 Month | Regulation of gap junction activity | 2 / 3 | 5.41 | 1.4e-05 |
|  | Scavenging by Class A Receptors | 2 / 7 | 5.66 | 6.4e-06 |
|  | Insertion of tail-anchored proteins into the endoplasmic reticulum membrane | 4 / 12 | 4.83 | 0.00015 |
|  | MTOR signaling | 5 / 14 | 3.71 | 0.006 |
|  | Norepinephrine Neurotransmitter Release Cycle | 8 / 15 | -4.10 | 0.0017 |
|  | Dopamine Neurotransmitter Release Cycle | 13 / 20 | -4.25 | 0.0015 |
|  | GABA synthesis, release, reuptake and degradation | 8 / 12 | -4.74 | 0.00018 |

Next, we examined pathways showing differential enrichment between the female and male datasets (enriched in proteins with high difference in fold changes). At the six-month time point, the only significant pathway that we identify is the “Interaction between L1 and Ankyrins pathways”. At the nine-month time point, we identified 11 differentially enriched pathways between females and males, all of which have previously been related to Alzheimer’s disease. These include several pathways related to apoptosis, cell cycle, axon guidance, and lipid homeostasis. In summary, our analysis reveals the sequential molecular signatures that drive AD progression. For additional significant pathways, please refer to Supplementary Data 6.

### 3.9. Validation on Independent Dataset: Confirming Phosphoproteomic Findings

To validate our findings and confirm the significance of candidate proteins, we analyzed an independent validation dataset for the 6-month and 9-month time points, each consisting of a smaller sample size (n = 4, 2 5XFAD and 2 control). In the 6-month dataset, we identified 2469 phosphopeptides in the main dataset and 3274 in the validation dataset, with 1330 phosphopeptides found to be common between the two. Using the same screening threshold and statistical analysis pipeline as described in previous sections (screening threshold: 5XFAD/WT 0.5 ≥ FC ≥ 2 and p ≤ 0.1), we investigated the significantly phosphorylated peptides. Among 12 peptides that passed the screening cutoffs in the main dataset and were present in the validation dataset, our analysis revealed 4 peptides that pass the same cutoffs in the validation dataset. These are: CLCN6-S726, GFAP-S12, GFAP-T40, and Stromal interaction molecule 2 (STIM2)-S28. Notably, the observed overlap of 4 peptides is statistically significant, as it exceeds the expected overlap of approximately 0.3 peptides, corresponding to an 11.7-fold increase compared to the expected overlap (Fisher p-value=2.4e-4).

At the 9-month time point, we identified 2576 phosphopeptides in the main dataset and 2806 in the validation dataset, with 1326 phosphopeptides found to be common between the two. Applying the same analysis, among 28 peptides that passed the cutoffs in the main dataset, 7 of them passed the cutoffs in the validation dataset as well: CLCN6-S774, GFAP-S12, A4-S441, PXDC2-S507, Ras-related protein Rab-3A (RAB3A)-S190, CaM kinase-like vesicle-associated protein (CAMKV)-T446, and Protein kinase C and casein kinase substrate in neurons protein 1 (PACN1)-S427. The observed overlap of these 7 peptides is statistically significant, exceeding the expected overlap of approximately 0.7 peptides, corresponding to a 10-fold increase compared to the expected overlap (Fisher p-value = 2.5e-6).

Next, we conducted a protein-level analysis of mean phosphorylation using the same thresholds (screening threshold: 5XFAD/WT 0.5 ≥ FC ≥ 2 and p ≤ 0.1). At the 6-month time point, out of the 1192 proteins in the main dataset and 1181 proteins in the validation dataset, 785 were common between the two. Among these, 7 proteins in the main dataset passed the cutoffs and 4 of them passed the cutoffs in both datasets: CLCN6, GFAP, APOE, and BIG3. Similarly, at the 9-month time point, we identified 1227 proteins in the main dataset and 1101 proteins in the validation dataset, with 788 proteins found to be common between the two. Among these, 11 proteins passed the cutoffs in the main dataset and 7 proteins in both: CLCN6, BIG3, GFAP, APOE, A4,

PXDC2, and DNJC5. Notably, at 6 months, we also observe that CLCN6 and STX7 passed the FDR cutoff of 0.1 in the validation dataset despite STX7 exhibiting fold changes less than 2. Similarly, at 9 months, GFAP and CLCN6 passed the FDR cutoff of 0.1. Overall, the validation results for candidate proteins are presented in Figure 5. For more detailed results, please refer to Supplementary Data 7.

**Figure 5.**
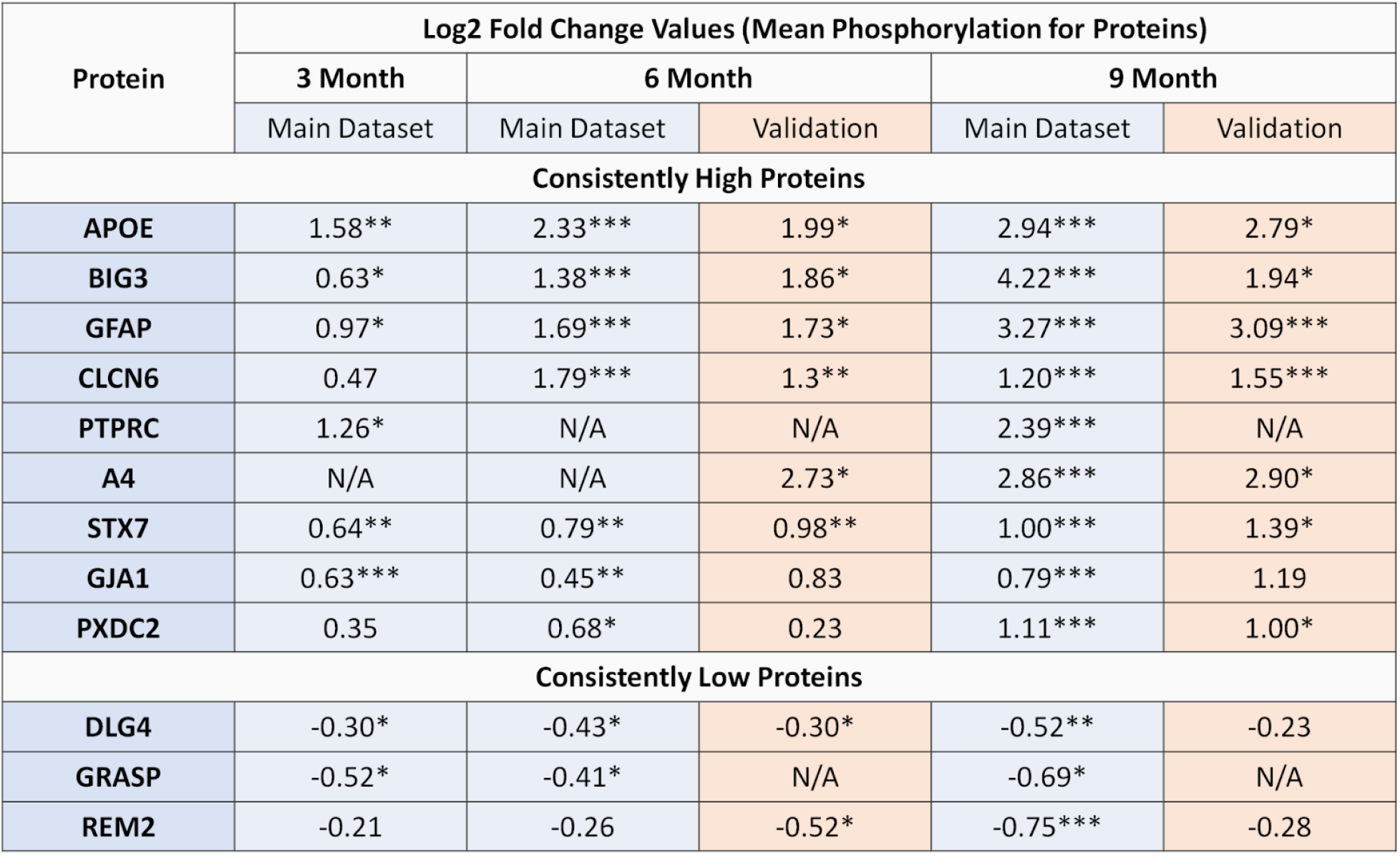
Validation data results for candidate proteins. Proteins are marked according to their statistical significance levels: * for p ≤ 0.1, ** for FDR ≤ 0.1 and *** for FDR ≤ 0.01.

## 4. Discussion

We started our analysis by investigating the interplay between protein expression and phosphorylation, which revealed only a minor association between the two omic datasets. This is on par with previously published data [Dammer, 2022] that found only modest correlation between different omic datasets in brain tissues, advocating for multi-level omics approaches to studying Alzheimer’s disease. Furthermore, we found that phosphorylation exhibited a 1.9x-4.4x higher percentage of peptides passing the screening criteria than protein expression across all timepoints. Remarkably, our sex-specific analysis revealed that females exhibit a substantially higher phosphorylation prevalence (4.4x) compared to protein expression in our earliest 3-month time point. Overall, these results suggest that dysregulation in protein phosphorylation is more prevalent and complementary to changes in protein expression, underscoring the importance of phosphorylation regulation in comprehending Alzheimer’s disease, particularly in its early stages.

Our analysis on the global phosphoproteomic differences revealed a balanced distribution of up- and down-regulated phosphopeptides in all timepoints, contrasting the proteomic data where more peptides were found to be upregulated than downregulated (greater than two thirds) [Lopes, 2022]. This dual regulation suggests that phosphorylation perpetuates the disease progression by disrupting homeostatic signals as much as it does by amplifying molecular signatures that exacerbate the disease. Surprisingly, a closer inspection revealed that the overlap among these phosphopeptides remained below 5% for all pairs of timepoints, implying distinct phosphorylation targeting at different AD stages. Additionally, our sex-specific analysis shows approximately 4.5x higher phospho-prevalence in females compared to males at 3 and 9 months, indicating that females exhibit higher phospho-dysregulation in both earlier and later stages of Alzheimer’s disease. This underscores the importance of developing sex-specific phosphorylation biomarkers that can be of potential clinical value for customized interventions in each sex.

Subsequently, we examined individual phosphosites displaying significant dysregulation at 3, 6, and 9 months (FDR ≤ 0.1; 0.5 ≥ FC ≥ 2). Remarkably, both temporal and sex-specific analyses consistently pinpointed the same top dysregulated proteins: APOE (S139, T140), BIG3 (S1646), CLCN6 (S726, S685, S774), GFAP (S12, Y13, T40) and STX7 (S45). In addition to the well-known proteins APOE and GFAP, we uncovered STX7 and CLCN6 as novel targets of phospho-dysregulation, suggesting AD’s involvement in the dysregulation of the late endocytic pathway [Jentsch, 2018]. Sex-specific analysis highlighted that the top dysregulated proteins in females are notably enriched in the Golgi apparatus. Moreover, enrichment analysis based on sexual differences revealed enriched GO terms encompassing endocytic recycling, neuron development, and neuron death.

Our biomarker analysis revealed several proteins that exhibit consistent phosphorylation signatures across the time points, including well-known hallmarks of Alzheimer’s disease such as APOE, GFAP, and A4. Furthermore, our analysis unveiled the biomarker potential of additional proteins, including BIG3, STX7, PTPRC, CLCN6, GJA1, PXDC2, and GRASP. These proteins play diverse biological roles. For example, STX7 is involved in the SNARE complex and plays a crucial role in the late endocytic pathway [Hankins, 2017; Sogorb-Esteve, 2022], while BIG3 is known to regulate GABA neurotransmitter release [Liu, 2016]. CLCN6, abundant in late endosomes of the nervous system, is found in amyloid plaques [Drummond, 2022] and associated with frontotemporal degeneration and dementia biomarkers [Sassi, 2021; Del Greco, 2011]. These findings suggest that CLCN6’s impact on autophagosome-mediated protein degradation might be critical in neurodegeneration.

Through our stoichiometry analysis, we examined the rate of phosphorylation among the top candidates identified in our biomarker analysis. Interestingly, we found that, while APOE and GFAP exhibit minimal phosphorylation rate (<1%) across all time points and samples, other candidates like STX7, GJA1, and BIG3 show a substantial rate of phosphorylation ranging from 20% to 40%. These contrasting profiles raise questions on the biological significance of the minor phosphorylation observed in APOE and GFAP, highlighting the potential relevance of the more pronounced phosphorylation events in STX7, GJA1, and BIG3. This prompts further exploration into the functional relevance of these phosphorylation events and their potential contributions to Alzheimer’s disease pathology.

Our kinase inference analysis indicated that PDK1 is significantly dysregulated, particularly at the 9 month time point in females. This observation is aligned with PDK1’s crucial involvement in synaptic plasticity and neuronal survival in the central nervous system (CNS), achieved by phosphorylating Akt, PKC, and p70S6, which are all pivotal for these processes [Bayascas, 2010]. In Alzheimer’s Disease, changes in PDK1 activity impact both the expression of Aβ and CDK5 through the PI3K/PDK1/Akt pathway [Pietri, 2013; Wen, 2008; Nava, 2011]. Our closer inspection into the PDK1 results at mRNA, protein expression and phosphorylation level indicated that PDK1 is a dysregulated kinase throughout different stages of Alzheimer’s Disease in the 5XFAD mouse model. PDK1 exhibits complex regulation at the protein expression and phosphorylation levels, which differ between male and females, with females exhibiting more pronounced changes. Whereas, the transcriptional regulation of PDK1 appears to be relatively stable over time. Further investigation into the downstream signaling pathways and targets of PDK1 may provide valuable insights into its role in Alzheimer’s Disease progression.

One of the significant findings in our Reactome pathway analysis was the regulation of the gap junction activity. The related gap junction alpha-1 protein, GJA1, was also a significant finding in our biomarker analysis, exhibiting consistent hyperphosphorylation across all time points. Gap junctions play a crucial role in the cell cycling, migration, and survival of neurons and glial cells [Zlomuzica, 2012]. Phosphorylation of gap junction proteins has been reported to regulate their assembly, trafficking, and stability [Lampe, 2010]. Whereas, another significant pathway that we identify, scavenging by class A receptors, has been associated with microglia’s ability to bind and phagocytize amyloid-beta (Aβ), and studies have shown a positive correlation between class A scavenger receptors and the accumulation of Aβ in the brain of transgenic AD mice [Bornemann, 2001].

Finally, AD-Xplorer, our online tool developed using the RokaiXplorer’s interactive data browser feature enables interactive exploration of our AD-related proteomics datasets and is accessible at https://yilmazs.shinyapps.io/ADXplorer. This platform enables researchers to investigate changes in phosphorylation, protein expression, and inferred kinase activities, complemented by a pathway enrichment analysis based on significant findings. It provides several interactive visualizations, including volcano plots, bar plots, box plots, heatmaps, and network views. The tool offers easy customization for focused subgroup analysis, such as selecting “9 Month” and “Female” to tailor the analysis to specific samples. To cater to diverse analytical needs, adjustable statistical cutoffs (based on p-values, fold changes or FDR) and various preprocessing options for quality control are provided. The tool supports exporting results as a formatted Excel report with detailed statistics and downloadable figure data. Overall, with an emphasis on temporal and sex-specific phosphoproteomic signatures, AD-Xplorer offers a user-friendly platform and facilitates in-depth exploration and interpretation of our dataset. It can aid in unraveling complexities of Alzheimer’s disease and serve as a valuable resource for biomarker discovery and therapeutic target identification.

## Abbreviations

A4: Amyloid-beta precursor protein
Aβ: Amyloid-beta
AD: Alzheimer’s disease
APOE: Apolipoprotein E (APOE)
BIG3: Brefeldin A-inhibited guanine nucleotide-exchange protein 3
CLCN6: Chloride Voltage-Gated Channel 6
DLG4: Disks large homolog 4
FC: Fold change
FDR: False discovery rate
GFAP: Glial fibrillary acidic protein
GJA1: Gap junction alpha-1 protein
GRASP: General receptor for phosphoinositides 1-associated scaffold protein
LC-MS: Liquid chromatography-tandem mass spectrometry
log2-FC: Log 2 transformed fold change
MS: Mass spectrometry
PDK1: Phosphoinositide-dependent kinase-1
PTMs: Post-translational modifications
PTPRC: Protein Tyrosine Phosphatase Receptor Type C
PXDC2: Plexin domain containing protein 2
REM2: Rad And Gem-Like GTP-Binding Protein 2
STX7: Syntaxin 7
WT: Wild type

## 5. Acknowledgments

This work was supported in part by the National Library of Medicine (NLM), National Institutes of General Medical Sciences (NIGMS), and the Office of the Director, National Institutes of Health (NIH), United States grants under R01-LM12980, R01-GM117208, R01-GM117208-03S1, S10-OD026882-01, and S10-OD028614-01. The content is solely the responsibility of the authors and does not necessarily represent the official views of the NLM, NIH or NIGMS.

## 6. Data availability

The data and code to replicate the analysis is deposited to Figshare and available with doi: 10.6084/m9.figshare.23903715. Additionally, the data and findings presented in this study can be explored interactively from AD-Xplorer application (https://yilmazs.shinyapps.io/ADXplorer/). This article contains supplemental data.

## 7. Author Contributions

DS, XQ, MK and MRC conceptualization; SY and FBTPL formal analysis; SY and FBTPL writing–original draft; SY, FBTPL, DS, XQ, MK and MRC writing–review and editing; FBTPL, DS, and RW investigation; SY and FBTPL data curation; SY and FBTPL software; SY visualization; SY, FBTPL, DS, XQ, MK and MRC methodology; SY, FBTPL and DS validation; XQ and MRC resources; MK and MRC supervision; MRC project administration; MK and MRC funding acquisition.

## Footnotes

1 http://explorer.rokai.io

2 https://yilmazs.shinyapps.io/ADXplorer

