## Supplementary Materials for "Exploring Temporal and Sex-Linked Dysregulation in Alzheimer’s Disease Phospho-Proteome"

#### Supplementary Methods

##### Statistical inference at protein level

To identify phosphopeptides that are significant, we perform moderated t-test, comparing the 5XFAD samples with the wildtype (WT) samples. Let $q[i]$ be the resultant log2 fold change, and $\sigma[i]$, $df[i]$ be the corresponding standard error and degrees of freedom obtained from moderated t-test for peptide $i$.

To identify potential biomarkers based on phosphorylation data that are consistent across time points, we perform the analysis at the protein level. For this purpose, we first compute the mean log-fold changes $q_p[j]$ for each protein $j$:

$$q_p[j] = \frac{\sum_{i \in \mathcal{V}_j} q[i]}{|\mathcal{V}_j|} \tag{1}$$

where $\mathcal{V}_j$ denotes the set of phosphopeptides corresponding to protein $j$.

To estimate the pooled standard error $\sigma_p[j]$ and the corresponding degrees of freedom $df_p[j]$ in the estimation of the mean log-fold changes for each protein $j$, we use the Satterthwaite approximation:

$$\begin{aligned} \sigma_p[j] &= \frac{\sqrt{\sum_{i \in \mathcal{V}_j} \sigma^2[i]}}{|\mathcal{V}_j|} \\ df_p[j] &= \frac{\left(\sum_{i \in \mathcal{V}_j} \sigma^2[i]\right)^2}{\sum_{i \in \mathcal{V}_j} \left(\frac{\sigma^4[i]}{df[i]}\right)} \end{aligned} \tag{2}$$

Based on these estimations, to compute the significance of a protein $j$, a t-test is performed with the t-statistic $t_p[j]$:

$$t_p[j] = \frac{q_p[j]}{\sigma_p[j]} \tag{3}$$

which follows a t-distribution with $df_p[j]$ degrees of freedom under the null hypothesis.

#### Consistency analysis to identify protein biomarkers

To assess consistency across time points, we introduce a simple statistic called the consistency score, which combines the log2-fold change results of individual time points. This score represents the total log2-fold change of proteins across time points, considering any missing values as log2-fold change of 0.

Let $\mathcal{T} = \{3 \text{ month}, 6 \text{ month}, 9 \text{ month}\}$ denote the set of time points, and let $q_p^{(t)}[j]$, $\sigma_p^{(t)}[j]$, and $df_p^{(t)}[j]$ respectively denote the log-fold changes, standard error, and the degrees of freedom for protein $j$ corresponding to the analysis performed for time point $t \in \mathcal{T}$. Based on these, we compute the consistency score $q_c[j]$ (i.e., total log fold change) for protein $j$ as follows:

$$q_c[j] = \sum_{t \in \mathcal{T}} q_p^{(t)}[j] \tag{4}$$

The corresponding standard error $\sigma_c[j]$ and the degrees of freedom $df_c[j]$ for the consistency score are estimated by the Satterthwaite approximation:

$$\begin{aligned} \sigma_c[j] &= \sqrt{\sum_{t \in \mathcal{T}} \left(\sigma_p^{(t)}[j]\right)^2} \\ df_p[j] &= \frac{\left(\sum_{t \in \mathcal{T}} \left(\sigma_p^{(t)}[j]\right)^2\right)^2}{\sum_{t \in \mathcal{T}} \left(\frac{\left(\sigma_p^{(t)}[i]\right)^4}{df_p^{(t)}[i]}\right)} \end{aligned} \tag{5}$$

Note that, if a protein $j$ have a missing data at time point $t$, we consider it to have $q_p^{(t)}[j] = 0$, $\sigma_p^{(t)}[j] = 0$ and $df_p^{(t)}[j] = \infty$ during the computation of the consistency score.

#### Statistical inference at pathway level

To understand the biological pathways and networks impacted by the observed phosphoproteome changes, we performed a quantitative pathway enrichment analysis based on the mean phosphorylation (log2-FC) of proteins.

Let $\mathcal{P}_k$ denote the set of proteins corresponding to pathway $k$ with non-missing data (i.e., each protein having at least one phosphopeptide identified in our dataset). To estimate the enrichment of pathway $k$, we compute the mean log fold change $q_e[k]$ for each protein in $\mathcal{P}_k$ as follows:

$$q_e[k] = \frac{\sum_{j \in \mathcal{P}_k} q_p[j]}{|\mathcal{P}_k|} \tag{6}$$

The corresponding standard error $\sigma_c[j]$ and the degrees of freedom $df_c[j]$ for the consistency score are estimated by the Satterthwaite approximation:

$$\sigma_e[k] = \frac{\sqrt{\sum_{j\in\mathcal{P}_k} \sigma_p^2[j]}}{|\mathcal{P}_k|}$$
$$df_e[k] = \frac{\left(\sum_{j\in\mathcal{P}_k} \sigma_p^2[j]\right)^2}{\sum_{j\in\mathcal{P}_k}\left(\frac{\sigma_p^4[j]}{df_p[j]}\right)} \tag{7}$$

Similar to the protein analysis, the statistical significance of a pathway $k$ is then assessed with a t-test, based on the t-statistic:

$$t_e[k] = \frac{q_e[k]}{\sigma_e[k]} \tag{8}$$

which follows a t-distribution with $df_e[k]$ degrees of freedom under the null hypothesis.

### Supplementary Figures

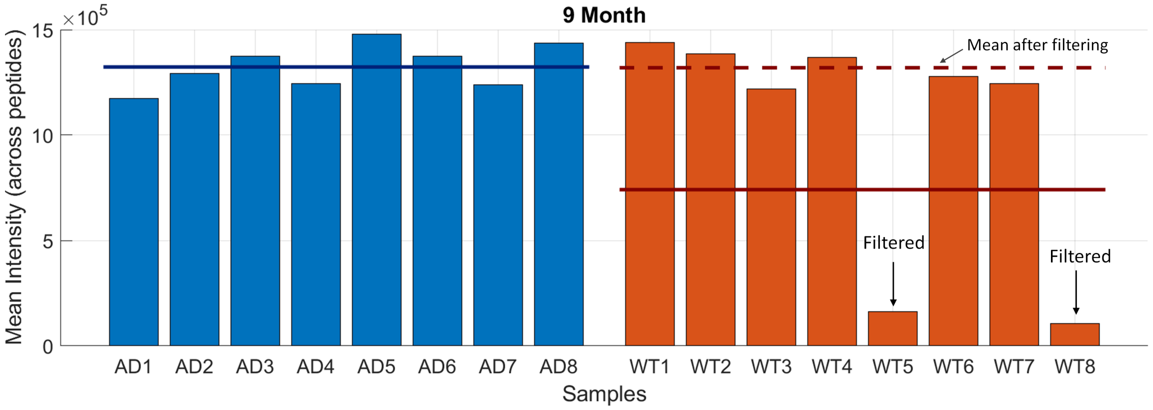


S. Figure 1: **Mean intensities across all phosphopeptides in the 9 month time point.** Samples named as AD1 to AD8 are in the 5XFAD group, and WT1 to WT8 are in the control (wildtype) group. Two WT samples with abnormally low intensities are marked on the plot. The solid lines indicate the mean values across samples. The dashed line on the WT group indicates the mean value after the two samples are filtered out.

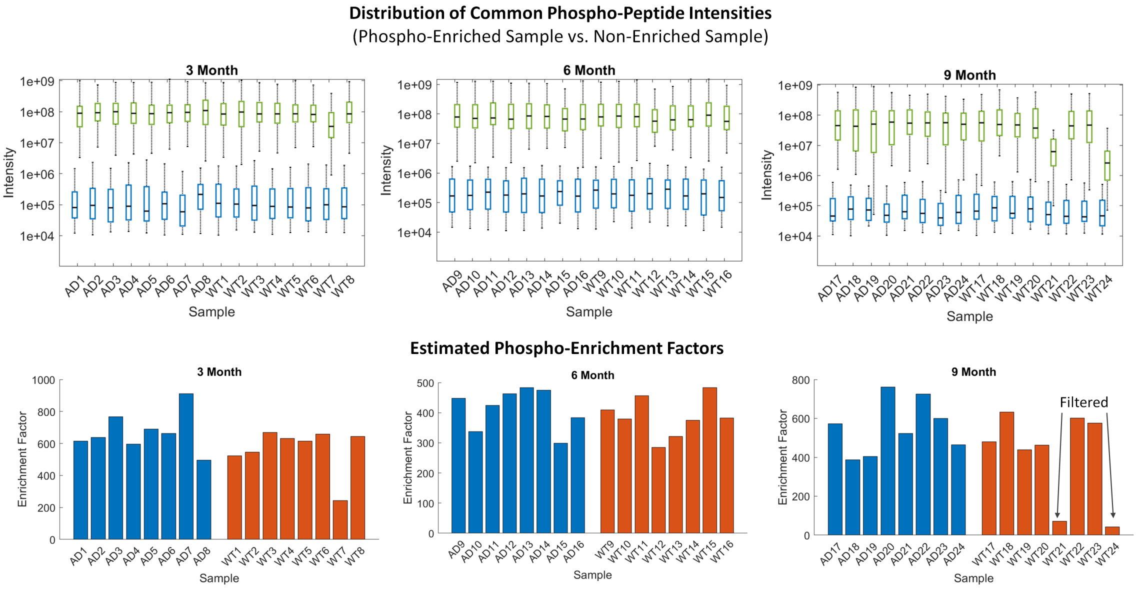


S. Figure 2: **Phospho-enriched vs. unenriched sample intensities.** (Top panels) Box plots indicating the distribution of the top phospho-peptide intensities for common proteins identified in phospho-enriched and unenriched samples. Each panel corresponds to a different time point. In each panel, green boxes on the top side correspond to phospho-enriched samples and blue boxes represent unenriched samples. (Bottom panels) Estimated phosphorylation enrichment factors based on the mean difference in the distribution of phospho-enriched samples and unenriched samples.

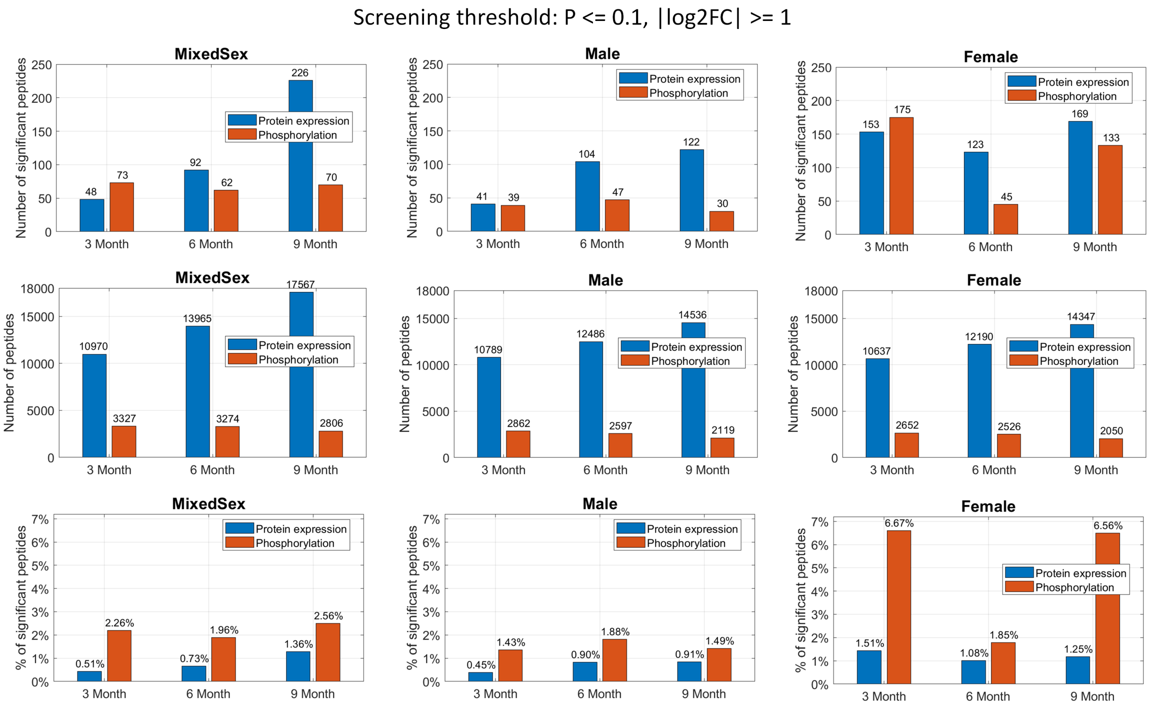


S. Figure 3: **Prevalence of phosphorylated or expressed peptides in Alzheimer's disease based on different time points and sex.** (Top panels) The number of peptides that pass the screening threshold ($p \leq 0.1$ and $0.5 \geq FC \geq 2$) for phosphorylation or expression for 3/6/9 month data. Left to right, each panel corresponds to a different analysis based on the sex of the samples: MixedSex, Male, and Female. (Middle panels) Total number of identified peptides. (Bottom panels) Percentage of the peptides that pass the screening threshold.

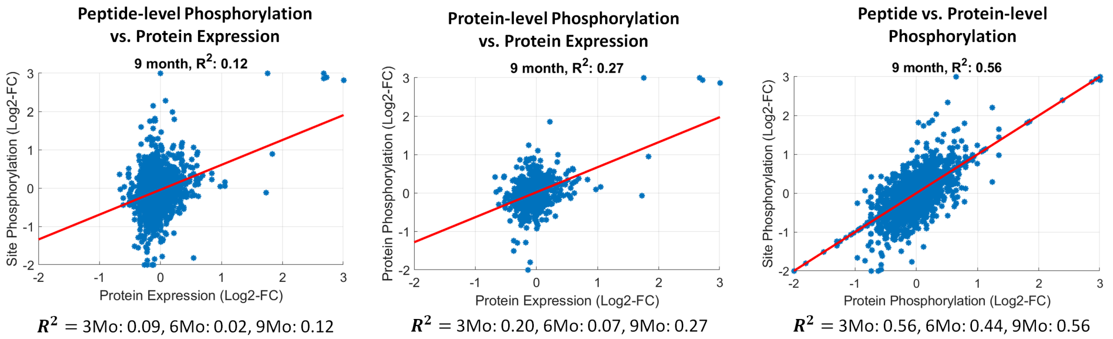


S. Figure 4: **Association and complementarity of phosphorylation information to protein expression in Alzheimer's disease phospho-proteome.** (Left) Scatter plot indicating the association between peptide-level phosphorylation and protein expression for the 9 month data. Each point represents a phospho-peptide and the x and y axes respectively indicate the log2 fold change values for protein expression and mean phosphorylation for the corresponding protein. (Left) Scatter plot indicating the association between phosphorylation and protein expression for the 9 month data. Each point represents a protein and the x and y axes respectively indicate log2 fold change values for protein expression and mean phosphorylation for the protein. (Right) Scatter plot indicating the association between phosphosite and protein-wise phosphorylation for the 9 month data. Each point represents a phosphosite and the x and y axes respectively indicate log2 fold change values for phosphorylation of the site and the mean phosphorylation for the corresponding protein. (All panels) The red line indicates the best fit line and the squared correlation ($R^2$) values for all timepoints are specified below the plots.

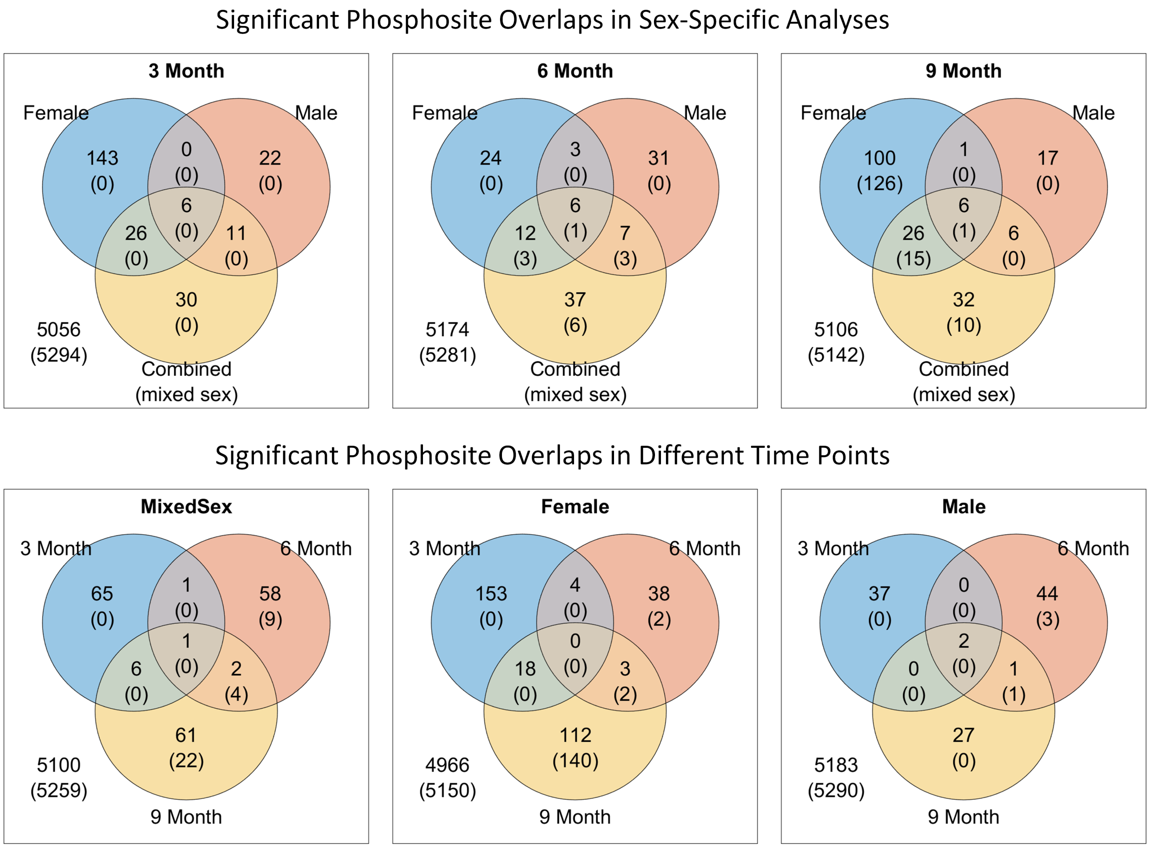


S. Figure 5: **Venn diagrams of the number of significant phosphosites for sex-specific analysis.** The numbers in the venn diagram indicate the number of phosphosites that pass the screening ($p \leq 0.1$ and $0.5 \geq FC \geq 2$). The numbers in parentheses indicate the number of significant phosphosites that pass $FDR \leq 0.1$. Top panels correspond to different time points (3/6/9 months), and bottom panels represent the type of sex-specific analysis (MixedSex/Female/Male).
